## Supplemental Figures S1-4 for "Post-ischemic ubiquitination at the postsynaptic density reversibly influences the activity of ischemia-relevant kinases"

### Supplementary Figures S1-4

accompanying the article "Post-ischemic ubiquitination at the postsynaptic density reversibly influences the activity of ischemia-relevant kinases."

Authors:
Luvna Dhawka[1, #], Victoria Palfini[1, #], Emma Hambright[1], Ismary Blanco[1], Carrie Poon[1], Anja Kahl[1], Ulrike Resch[2], Ruchika Bhawal[3], Corinne Benakis[1, &], Vaishali Balachandran[1], Sheng Zhang[3], Costantino Iadecola[1], Karin Hochrainer[1, *]

[1]Feil Family Brain and Mind Research Institute, Weill Cornell Medicine, New York, NY10065, USA; [2]Center for Physiology and Pharmacology, Medical University of Vienna, 1090 Vienna, Austria; [3]Institute of Biotechnology, Cornell University, Ithaca, NY14850, USA.

[&]Current address: Institute for Stroke and Dementia Research, Ludwig-Maximilians-University Munich, 81377 Munich, Germany.
[#]These authors contributed equally to this work.

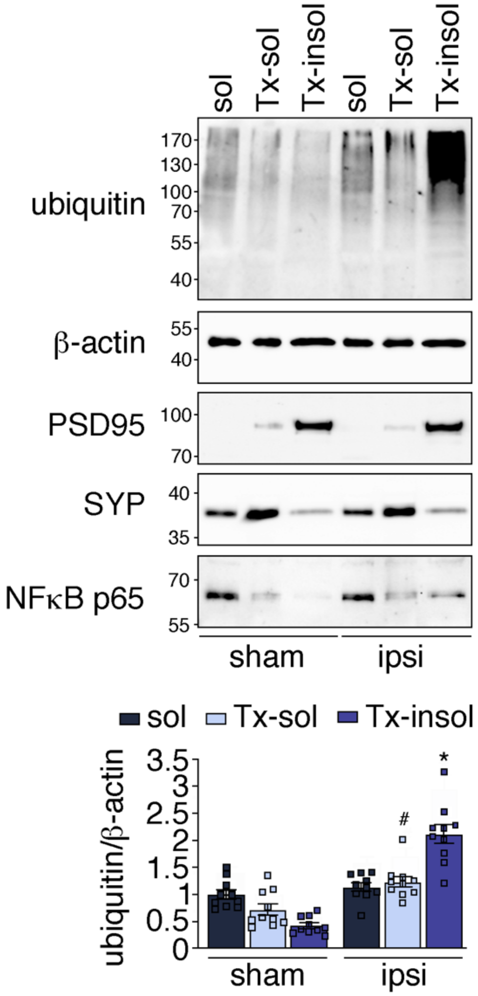


Figure S1. **The ubiquitin-containing Tx-insoluble fraction stains positive for the postsynaptic density marker PSD95.** Ubiquitin was detected in cortical soluble (sol), Tx-soluble (Tx-sol) and Tx-insoluble (Tx-insol) fractions derived from mice that underwent sham or MCAO/1h reperfusion (ipsi) surgeries. The Tx-insol fraction was also stained with PSD95, showing that this fraction contains the postsynaptic density. Markers for cytosol (p65) and synaptic membranes (SYP) were found in sol and Tx-sol fractions, respectively. Results were quantified. n=10 mice/group; #P=0.0023 from Tx-sol sham; *P<0.0001 from Tx-insol sham.

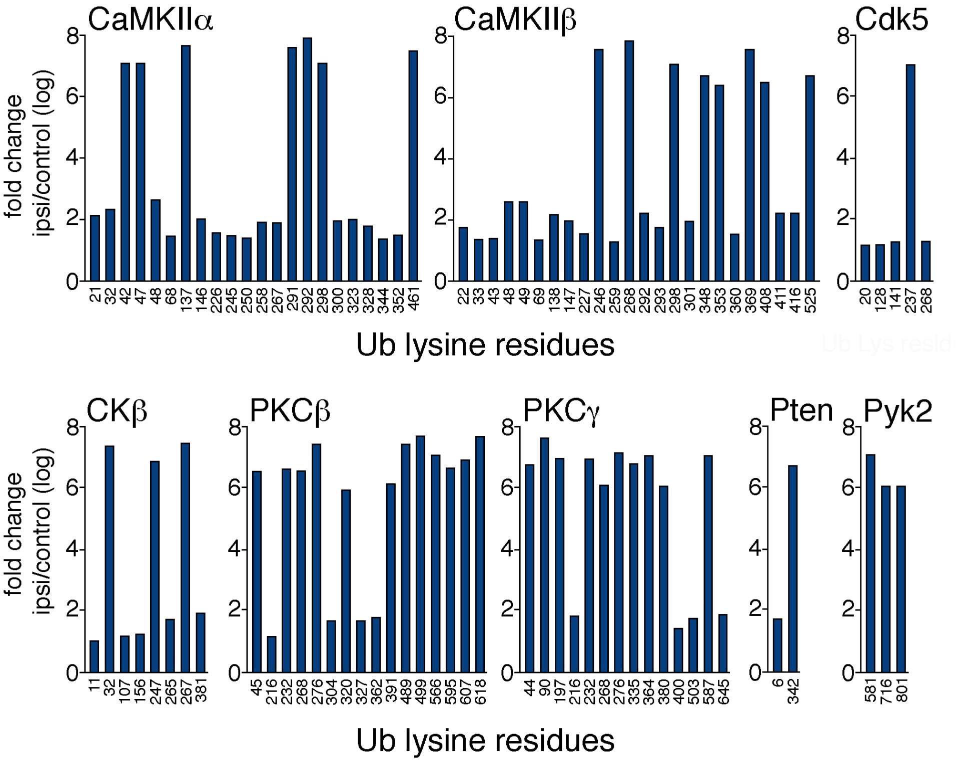


Figure S2. **Fold increase of ubiquitination on lysine residues in PSD-associated kinases and phosphatases after MCAO.** Ubiquitinated lysine residues in the kinases CaMKIIα, CaMKIIβ, PKCβ, PKCγ, Cdk5, CKβ, and Pyk2, and the phosphatase Pten as identified by nanoLC-MS/MS. Mean fold change ipsilateral vs control cortex (pooled sham and contralateral) is shown on a $\log_{10}$ scale (n=2-3 MS runs). Ub, ubiquitin.

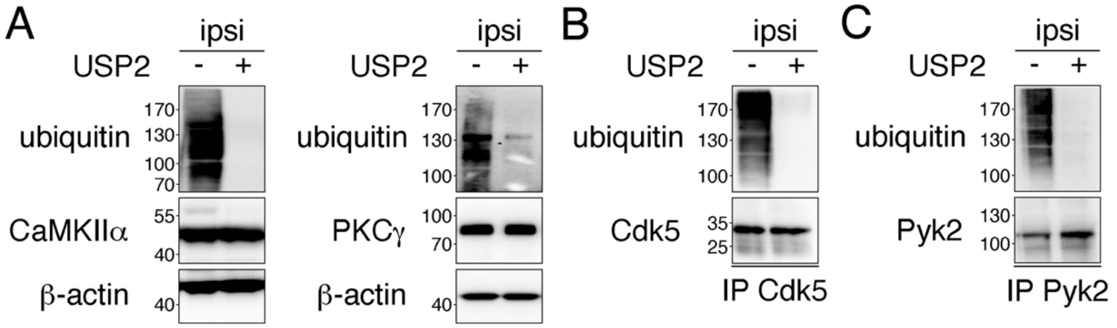


Figure S3. **USP2 efficiently removes ubiquitin from substrates.** Total PSD lysates or precipitated PSD-kinases (for Cdk5 and Pyk2) were incubated with recombinant USP2, and high molecular weight ubiquitin smear was detected by Western Blot. **(A)** CaMKII and PKC, as well as **(B)** Cdk5 and **(C)** Pyk2 kinases and β-actin were used as loading controls. ipsi, ipsilateral.

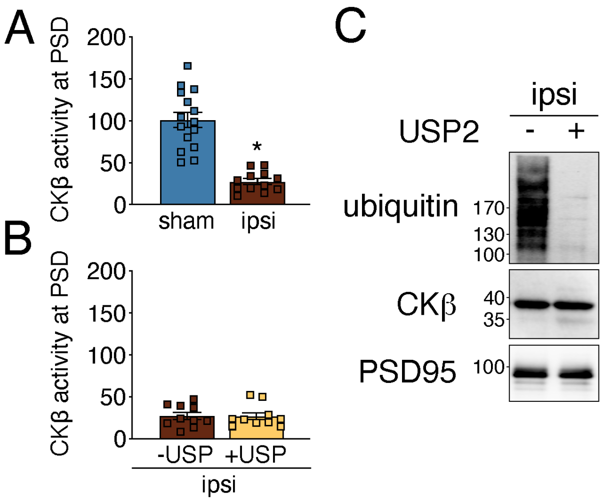


Figure S4. **The effect of MCAO/reperfusion and USP2 treatment on CKβ activity. (A)** CKβ activity was determined in cortical PSD lysates from sham and stroked mice. *P<0.0001, n=12-15 animals/group. **(B)** PSD lysates from stroked mice were treated with USP2 and CKβ activity was reassessed. Not significant, n=10 animals. **(C)** USP2 digest was verified by Western Blotting as in described in Figure S3. CKβ and PSD95 were loaded as controls. ipsi, ipsilateral.
