## Supplemental Tables S2-4 for "Post-ischemic ubiquitination at the postsynaptic density reversibly influences the activity of ischemia-relevant kinases"

### Supplementary Tables S2-4

accompanying the article “Post-ischemic ubiquitination at the postsynaptic density reversibly influences the activity of ischemia-relevant kinases.”

Authors:
Luvna Dhawka[1, #], Victoria Palfini[1, #], Emma Hambright[1], Ismary Blanco[1], Carrie Poon[1], Anja Kahl[1], Ulrike Resch[2], Ruchika Bhawal[3], Corinne Benakis[1, &], Vaishali Balachandran[1], Sheng Zhang[3], Costantino Iadecola[1], Karin Hochrainer[1, *]

[1]Feil Family Brain and Mind Research Institute, Weill Cornell Medicine, New York, NY10065, USA; [2]Center for Physiology and Pharmacology, Medical University of Vienna, 1090 Vienna, Austria; [3]Institute of Biotechnology, Cornell University, Ithaca, NY14850, USA.

[&]Current address: Institute for Stroke and Dementia Research, Ludwig-Maximilians-University Munich, 81377 Munich, Germany.
[#]These authors contributed equally to this work.

Table S2: **Functional annotation of proteins with increased post-ischemic ubiquitination (in alphabetical order).**

| Gene name and Association | Description | Entrez Gene ID |
|---|---|---|
| **cytoskeletal proteins** | | |
| Cep170b | centrosomal protein 170B | 217882 |
| Dynll1 | dynein light chain LC8-type 1 | 56455 |
| Elmo2 | engulfment and cell motility 2 | 140579 |
| Epb4.1l1 | erythrocyte membrane protein band 4.1 like 1 | 13821 |
| Map1b | microtubule-associated protein 1B | 17755 |
| Mapre2 | microtubule-associated protein, RP/EB family, member 2 | 212307 |
| Phactr1 | phosphatase and actin regulator 1 | 218194 |
| Syne1 | spectrin repeat containing, nuclear envelope 1 | 64009 |
| Tubb2a | tubulin, beta 2A class IIA | 22151 |
| Tubb4b | tubulin, beta 4B class IVB | 227613 |
| Tubb5 | tubulin, beta 5 class I | 22154 |
| **G-proteins /GTP regulators** | | |
| Abr | active BCR-related gene | 109934 |
| Agap2 | ArfGAP with GTPase domain, ankyrin repeat and PH domain 2 | 216439 |
| Arfgap1 | ADP-ribosylation factor GTPase activating protein 1 | 228998 |
| Arhgef2 | rho/rac guanine nucleotide exchange factor (GEF) 2 | 16800 |
| Bcr | BCR activator of RhoGEF and GTPase | 110279 |
| Farp1 | FERM, RhoGEF (Arhgef) and pleckstrin domain protein 1 (chondrocyte-derived) | 223254 |
| Gnai1 | guanine nucleotide binding protein (G protein), alpha inhibiting 1 | 14677 |
| Gnai2 | guanine nucleotide binding protein (G protein), alpha inhibiting 2 | 14678 |
| Gnai3 | guanine nucleotide binding protein (G protein), alpha inhibiting 3 | 14679 |
| Gnal | guanine nucleotide binding protein, alpha stimulating, olfactory type | 14680 |
| Gnao1 | guanine nucleotide binding protein, alpha O | 14681 |
| Gnas | GNAS (guanine nucleotide binding protein, alpha stimulating) complex locus | 14683 |
| Gnat1 | guanine nucleotide binding protein, alpha transducing 1 | 14685 |
| Gnat2 | guanine nucleotide binding protein, alpha transducing 2 | 14686 |

| | | |
|---|---|---|
| Gnat3 | guanine nucleotide binding protein, alpha transducing 3 | 242851 |
| Gpsm1 | G-protein signalling modulator 1 (AGS3-like, C. elegans) | 67839 |
| Iqgap2 | IQ motif containing GTPase activating protein 2 | 544963 |
| Iqsec1 | IQ motif and Sec7 domain 1 | 232227 |
| Iqsec2 | IQ motif and Sec7 domain 2 | 245666 |
| Kalrn | kalirin, RhoGEF kinase | 545156 |
| Psd | pleckstrin and Sec7 domain containing | 73728 |
| Rasal2 | RAS protein activator like 2 | 226525 |
| Rasgrf1 | RAS protein-specific guanine nucleotide-releasing factor 1 | 19417 |
| Rasgrp2 | RAS, guanyl releasing protein 2 | 19395 |
| Syngap1 | synaptic Ras GTPase activating protein 1 homolog (rat) | 240057 |
| **kinases /phosphatases** | | |
| Ak1 | adenylate kinase 1 | 11636 |
| Ak5 | adenylate kinase 5 | 229949 |
| Brsk1 | BR serine/threonine kinase 1 | 381979 |
| Camk1d | calcium/calmodulin-dependent protein kinase ID | 227541 |
| Camk2a | calcium/calmodulin-dependent protein kinase II alpha | 12322 |
| Camk2b | calcium/calmodulin-dependent protein kinase II, beta | 12323 |
| Camk2d | calcium/calmodulin-dependent protein kinase II, delta | 108058 |
| Camk2g | calcium/calmodulin-dependent protein kinase II gamma | 12325 |
| Camk4 | calcium/calmodulin-dependent protein kinase IV | 12326 |
| Camkk1 | calcium/calmodulin-dependent protein kinase kinase 1, alpha | 55984 |
| Camkv | CaM kinase-like vesicle-associated | 235604 |
| Cdk16 | cyclin-dependent kinase 16 | 18555 |
| Cdk17 | cyclin-dependent kinase 17 | 237459 |
| Cdk18 | cyclin-dependent kinase 18 | 18557 |
| Cdk5 | cyclin-dependent kinase 5 | 12568 |
| Ckb | creatine kinase, brain | 12709 |
| Dgkb | diacylglycerol kinase, beta | 217480 |
| Dgkh | diacylglycerol kinase, eta | 380921 |
| Dusp3 | dual specificity phosphatase 3 (vaccinia virus phosphatase VH1-related) | 72349 |

| | | |
|---|---|---|
| Itpka | inositol 1,4,5-trisphosphate 3-kinase A | 228550 |
| Itpkb | inositol 1,4,5-trisphosphate 3-kinase B | 320404 |
| Map4k3 | mitogen-activated protein kinase kinase kinase kinase 3 | 225028 |
| Mast3 | microtubule associated serine/threonine kinase 3 | 546071 |
| Mtmr6 | myotubularin related protein 6 | 219135 |
| Ntrk2 | neurotrophic tyrosine kinase, receptor, type 2 | 18212 |
| Pnck | pregnancy upregulated non-ubiquitously expressed CaM kinase | 93843 |
| Ppp2cb | protein phosphatase 2 (formerly 2A), catalytic subunit, beta isoform | 19053 |
| Prkca | protein kinase C, alpha | 18750 |
| Prkcb | protein kinase C, beta | 18751 |
| Prkcg | protein kinase C, gamma | 18752 |
| Pten | phosphatase and tensin homolog | 19211 |
| Ptk2b | PTK2 protein tyrosine kinase 2 beta | 19229 |
| Stk39 | serine/threonine kinase 39 | 53416 |
| **metabolic enzymes** | | |
| Abhd6 | abhydrolase domain containing 6 | 66082 |
| Cbr1 | carbonyl reductase 1 | 12408 |
| Cyp46a1 | cytochrome P450, family 46, subfamily a, polypeptide 1 | 13116 |
| Gapdh | glyceraldehyde-3-phosphate dehydrogenase | 14433 |
| Gsto1 | glutathione S-transferase omega 1 | 14873 |
| Hdlbp | high density lipoprotein (HDL) binding protein | 110611 |
| Nmral1 | NmrA-like family domain containing 1 | 67824 |
| Pgam2 | phosphoglycerate mutase 2 | 56012 |
| Srr | serine racemase | 27364 |
| Tpi1 | triosephosphate isomerase 1 | 21991 |
| **other proteins /proteins with unknown function** | | |
| Abcf2 | ATP-binding cassette, sub-family F (GCN20), member 2 | 27407 |
| Anks1b | ankyrin repeat and sterile alpha motif domain containing 1B | 77531 |
| Armc6 | armadillo repeat containing 6 | 76813 |
| Cpne6 | copine VI | 12891 |
| Dmwd | dystrophia myotonica-containing WD repeat motif | 13401 |

| | | |
|---|---|---|
| Etl4 | enhancer trap locus 4 | 208618 |
| Fam81a | family with sequence similarity 81, member A | 76886 |
| Gm15294 | predicted gene 15294 | 666488 |
| Gm3839 | predicted pseudogene 3839 | 100042427 |
| Gm49325 | predicted gene, 49325 | N/A |
| Gm8797 | predicted pseudogene 8797 | 667759 |
| Hpca | hippocalcin | 15444 |
| Lzts1 | leucine zipper, putative tumor suppressor 1 | 211134 |
| Mpp1 | membrane protein, palmitoylated | 17524 |
| Mtfr1l | mitochondrial fission regulator 1-like | 76824 |
| Nap1l1 | nucleosome assembly protein 1-like 1 | 53605 |
| Phf24 | PHD finger protein 24 | 230085 |
| Pin1rt1 | protein (peptidyl-prolyl cis/trans isomerase) NIMA-interacting 1, retrogene 1 | 241593 |
| Prickle2 | prickle planar cell polarity protein 2 | 243548 |
| Rufy3 | RUN and FYVE domain containing 3 | 52822 |
| Zwint | ZW10 interactor | 52696 |
| **other signaling proteins** | | |
| Cnksr2 | connector enhancer of kinase suppressor of Ras 2 | 245684 |
| Cnrip1 | cannabinoid receptor interacting protein 1 | 380686 |
| Hpcal1 | hippocalcin-like 1 | 53602 |
| Hpcal4 | hippocalcin-like 4 | 170638 |
| Nyap1 | neuronal tyrosine-phosphorylated phosphoinositide 3-kinase adaptor 1 | 243300 |
| Pea15a | phosphoprotein enriched in astrocytes 15A | 18611 |
| Srcin1 | SRC kinase signaling inhibitor 1 | 56013 |
| Tollip | toll interacting protein | 54473 |
| **proteins regulating protein homeostasis** | | |
| Anp32e | acidic (leucine-rich) nuclear phosphoprotein 32 family, member E | 66471 |
| Cul9 | cullin 9 | 78309 |
| Dnajc7 | DnaJ heat shock protein family (Hsp40) member C7 | 56354 |
| Fkbp8 | FK506 binding protein 8 | 14232 |
| Hecw1 | HECT, C2 and WW domain containing E3 ubiquitin protein ligase 1 | 94253 |

| | | |
|---|---|---|
| Hspa12a | heat shock protein 12A | 73442 |
| Hspa4l | heat shock protein 4 like | 18415 |
| Kxd1 | KxDL motif containing 1 | 75620 |
| Lrsam1 | leucine rich repeat and sterile alpha motif containing 1 | 227738 |
| Nub1 | negative regulator of ubiquitin-like proteins 1 | 53312 |
| Nudcd2 | NudC domain containing 2 | 52653 |
| Nudcd3 | NudC domain containing 3 | 209586 |
| Rps27a | ribosomal protein S27A | 78294 |
| Sumo2 | small ubiquitin-like modifier 2 | 170930 |
| Sumo3 | small ubiquitin-like modifier 3 | 20610 |
| Trim28 | tripartite motif-containing 28 | 21849 |
| Trim9 | tripartite motif-containing 9 | 94090 |
| Uba52 | ubiquitin A-52 residue ribosomal protein fusion product 1 | 22186 |
| Ubb | ubiquitin B | 22187 |
| Ubc | ubiquitin C | 22190 |
| Ube2v1 | ubiquitin-conjugating enzyme E2 variant 1 | 66589 |
| Ube3c | ubiquitin protein ligase E3C | 100763 |
| Usp13 | ubiquitin specific peptidase 13 (isopeptidase T-3) | 72607 |
| Usp5 | ubiquitin specific peptidase 5 (isopeptidase T) | 22225 |
| Vcpip1 | valosin containing protein (p97)/p47 complex interacting protein 1 | 70675 |
| **receptors /channels** | | |
| Adgrb1 | adhesion G protein-coupled receptor B1 | 107831 |
| Adgrb3 | adhesion G protein-coupled receptor B3 | 210933 |
| Gria2 | glutamate receptor, ionotropic, AMPA2 (alpha 2) | 14800 |
| Gria3 | glutamate receptor, ionotropic, AMPA3 (alpha 3) | 53623 |
| Grin1 | glutamate receptor, ionotropic, NMDA1 (zeta 1) | 14810 |
| Grin2a | glutamate receptor, ionotropic, NMDA2A (epsilon 1) | 14811 |
| Grin2b | glutamate receptor, ionotropic, NMDA2B (epsilon 2) | 14812 |
| Nr3c1 | nuclear receptor subfamily 3, group C, member 1 | 14815 |
| **proteins involved in RNA and protein synthesis** | | |
| Abce1 | ATP-binding cassette, sub-family E (OABP), member 1 | 24015 |

| | | |
|---|---|---|
| Apbb1 | amyloid beta (A4) precursor protein-binding, family B, member 1 | 11785 |
| Coil | coilin | 12812 |
| Csde1 | cold shock domain containing E1, RNA binding | 229663 |
| Ddx5 | DEAD box helicase 5 | 13207 |
| Hnrnpa0 | heterogeneous nuclear ribonucleoprotein A0 | 77134 |
| Hnrnpa1 | heterogeneous nuclear ribonucleoprotein A1 | 15382 |
| Hnrnpc | heterogeneous nuclear ribonucleoprotein C | 15381 |
| Hnrnpf | heterogeneous nuclear ribonucleoprotein F | 98758 |
| Hnrnph1 | heterogeneous nuclear ribonucleoprotein H1 | 59013 |
| Hnrnph2 | heterogeneous nuclear ribonucleoprotein H2 | 56258 |
| Hnrnpm | heterogeneous nuclear ribonucleoprotein M | 76936 |
| Hnrnpu | heterogeneous nuclear ribonucleoprotein U | 51810 |
| Hnrnpul2 | heterogeneous nuclear ribonucleoprotein U-like 2 | 68693 |
| Matr3 | matrin 3 | 17184 |
| Mrtfb | myocardin related transcription factor B | 239719 |
| Nab2 | Ngfi-A binding protein 2 | 17937 |
| Nacc1 | nucleus accumbens associated 1, BEN and BTB (POZ) domain containing | 66830 |
| Pabpc4 | poly(A) binding protein, cytoplasmic 4 | 230721 |
| Pcif1 | phosphorylated CTD interacting factor 1 | 228866 |
| Pelo | pelota mRNA surveillance and ribosome rescue factor | 105083 |
| Prpf3 | pre-mRNA processing factor 3 | 70767 |
| Purb | purine rich element binding protein B | 19291 |
| Rtca | RNA 3'-terminal phosphate cyclase | 66368 |
| Tardbp | TAR DNA binding protein | 230908 |
| **scaffolding /clustering proteins** | | |
| Begain | brain-enriched guanylate kinase-associated | 380785 |
| Caskin1 | CASK interacting protein 1 | 268932 |
| Dlg1 | discs large MAGUK scaffold protein 1 | 13383 |
| Dlg2 | discs large MAGUK scaffold protein 2 | 23859 |
| Dlg3 | discs large MAGUK scaffold protein 3 | 53310 |
| Dlg4 | discs large MAGUK scaffold protein 4 | 13385 |

| | | |
|---|---|---|
| Dlgap1 | DLG associated protein 1 | 224997 |
| Dlgap2 | DLG associated protein 2 | 244310 |
| Dlgap3 | DLG associated protein 3 | 242667 |
| Dlgap4 | DLG associated protein 4 | 228836 |
| Lrrc7 | leucine rich repeat containing 7 | 242274 |
| Mpp2 | membrane protein, palmitoylated 2 (MAGUK p55 subfamily member 2) | 50997 |
| Pclo | piccolo (presynaptic cytomatrix protein) | 26875 |
| Shank1 | SH3 and multiple ankyrin repeat domains 1 | 243961 |
| Shank2 | SH3 and multiple ankyrin repeat domains 2 | 210274 |
| Sorbs2 | sorbin and SH3 domain containing 2 | 234214 |
| **trafficking /transport proteins** | | |
| Apba2 | amyloid beta (A4) precursor protein-binding, family A, member 2 | 11784 |
| Clvs2 | clavesin 2 | 215890 |
| Grasp | trafficking regulator and scaffold protein tamalin | 56149 |
| Gripap1 | GRIP1 associated protein 1 | 54645 |
| Lin7a | lin-7 homolog A (C. elegans) | 108030 |
| Napa | N-ethylmaleimide sensitive fusion protein attachment protein alpha | 108124 |
| Napb | N-ethylmaleimide sensitive fusion protein attachment protein beta | 17957 |
| Nsf | N-ethylmaleimide sensitive fusion protein | 18195 |
| Rab11fip5 | RAB11 family interacting protein 5 (class I) | 52055 |
| Snap47 | synaptosomal-associated protein, 47 | 67826 |
| Snx3 | sorting nexin 3 | 54198 |
| Stam | signal transducing adaptor molecule (SH3 domain and ITAM motif) 1 | 20844 |
| Unc13a | unc-13 homolog A | 382018 |
| Unc13b | unc-13 homolog B | 22249 |
| Unc13c | unc-13 homolog C | 208898 |
| Vps37a | vacuolar protein sorting 37A | 52348 |

Table S3: **Primary antibodies used for immunoprecipitation.**

| Antibody | Clone | Host | IgG subclass | Dilution | Manufacturer | Catalog number |
|---|---|---|---|---|---|---|
| CaMKIIα | CBα-2 | Mouse | $IgG_{2a}$ | 4μg | Invitrogen | 13-7300 |
| CaMKIIβ | CB-β-I | Mouse | $IgG_{2b}$ | 4μg | Invitrogen | 13-9800 |
| Cdk5 | -- | Rabbit | -- | 1:50 | Cell Signaling | 2506 |
| CKβ | -- | Rabbit | -- | 4μg | Proteintech | 15137-1-AP |
| GluA2 | E1L8U | Rabbit | -- | 1:50 | Cell Signaling | 13607 |
| GluN1 | D65B7 | Rabbit | -- | 1μl/50μg | Cell Signaling | 5704 |
| GluN2B | 13/NMDAR2B | Mouse | $IgG_{2b}$ | 4μg | BD Biosciences | 610416 |
| p35/p25 | C64B10 | Rabbit | -- | 1:50 | Cell Signaling | 2680 |
| PKCβ | D3E70 | Rabbit | -- | 1:50 | Cell Signaling | 46809 |
| PKCγ | D2V6T | Rabbit | -- | 1:100 | Cell Signaling | 59090 |
| PSD93 | D4Z4D | Rabbit | -- | 1:50 | Cell Signaling | 19046 |
| PSD95 | 6G6-1C9 | Mouse | $IgG_{2a}$ | 4μg | Invitrogen | MA1-045 |
| Pten | -- | Rabbit | -- | 2μg | Proteintech | 22034-1-AP |
| Pyk2 | -- | Rabbit | -- | 4μg | Proteintech | 17592-1-AP |
| Shank2 | -- | Rabbit | -- | 1:50 | Cell Signaling | 12218 |
| Shank3 | D5K6R | Rabbit | -- | 1μl/50μg | Cell Signaling | 64555 |
| TrkB | -- | Rabbit | -- | 1:200 | Proteintech | 13129-1-AP |
| Isotype control | G3A1 | Mouse | $IgG_1$ | 4μg | Cell Signaling | 5415 |
| Isotype control | MOPC-173 | Mouse | $IgG_{2a}$ | 4μg | Biolegend | 400201 |
| Isotype control | 27-35 | Mouse | $IgG_{2b}$ | 4μg | Biolegend | 402201 |
| Isotype control | poly29108 | Rabbit | -- | 4μg | Biolegend | 910801 |

Note: For Cell Signaling antibodies, dilutions are μL antibody per μg total protein

Table S4: **Primary antibodies used for Western blotting.**

| Antibody | Clone | Host | IgG subclass | Dilution | Manufacturer | Catalog number |
|---|---|---|---|---|---|---|
| β-actin | AC-15 | Mouse | $IgG_1$ | 1:10000 | Sigma | A5441 |
| CaMKIIα | CBα-2 | Mouse | $IgG_{2a}$ | 1:500 | Invitrogen | 13-7300 |
| CaMKIIβ | CB-β-I | Mouse | $IgG_{2b}$ | 1:1000 | Invitrogen | 13-9800 |
| Cdk5 | -- | Rabbit | -- | 1:1000 | Cell Signaling | 2506 |
| CKβ | -- | Rabbit | -- | 1:1000 | Proteintech | 15137-1-AP |
| Crmp2 | C terminus | Rabbit | -- | 1:1000 | ECM Biosciences | CP2161 |
| nNOS | -- | Rabbit | -- | 1:2000 | Enzo | BML-SA227 |
| GluA2 | E1L8U | Rabbit | -- | 1:1000 | Cell Signaling | 13607 |
| GluN1 | D65B7 | Rabbit | -- | 1:1000 | Cell Signaling | 5704 |
| GluN2B | 13/NMDAR2B | Mouse | $IgG_{2b}$ | 1:500 | BD Biosciences | 610416 |
| p35/p25 | C64B10 | Rabbit | -- | 1:1000 | Cell Signaling | 2680 |
| PKCβ | D3E70 | Rabbit | -- | 1:1000 | Cell Signaling | 46809 |
| PKCγ | D2V6T | Rabbit | -- | 1:1000 | Cell Signaling | 59090 |
| PSD93 | D4Z4D | Rabbit | -- | 1:1000 | Cell Signaling | 19046 |
| PSD95 | 16/PSD-95 | Mouse | $IgG_1$ | 1:500 | BD Biosciences | 610495 |
| Pten | -- | Rabbit | -- | 1:2000 | Proteintech | 22034-1-AP |
| Pyk2 | -- | Rabbit | -- | 1:3000 | Proteintech | 17592-1-AP |
| Shank2 | -- | Rabbit | -- | 1:1000 | Cell Signaling | 12218 |
| Shank3 | D5K6R | Rabbit | -- | 1:1000 | Cell Signaling | 64555 |
| Src | 36D10 | Rabbit | -- | 1:1000 | Cell Signaling | 2109 |
| Tau | TAU-5 | Mouse | $IgG_1$ | 1:200 | Invitrogen | MA5-12808 |
| TrkB | -- | Rabbit | -- | 1:1000 | Proteintech | 13129-1-AP |
| Ubiquitin | Ubi-1 | Mouse | $IgG_1$ | 1:500 | Invitrogen | 13-1600 |
| Phospho-Crmp2 (S522) | -- | Rabbit | -- | 1:1000 | ECM Biosciences | CP2191 |
| Phospho-GluN2B (Y1472) | -- | Rabbit | -- | 1:1000 | Cayman Chemical | 10009761 |
| Phospho-GluN2B (S1303) | -- | Rabbit | -- | 1μg/mL | Millipore | 07-398 |
| Phospho-nNOS (S847) | -- | Rabbit | -- | 1μg/mL | Abcam | ab16650 |
| Phospho-Pyk2 (Y402) | -- | Rabbit | -- | 1:1000 | Invitrogen | 44-618G |
| Phospho-serine/threonine | -- | Rabbit | -- | 1:1000 | Cell Signaling | 9631 |
| Phospho-Src (Y419) | D49G4 | Rabbit | -- | 1:1000 | Cell Signaling | 6943 |
| Phospho-Tau (Y18) | 9G3 | Mouse | $IgG_{2a}$ | 1:1000 | Novus Biologicals | NBP2-42402 |
| Phospho-Tau (S202/205) | AT8 | Mouse | $IgG_1$ | 1:500 | Invitrogen | MN1020 |
| Phospho-tyrosine | P-Tyr-100 | Mouse | $IgG_1$ | 1:2000 | Cell Signaling | 9411 |
